# A theory and recipe to construct general and biologically plausible integrating continuous attractor neural networks

**DOI:** 10.1101/2025.05.07.652608

**Authors:** Federico Claudi, Sarthak Chandra, Ila R. Fiete

## Abstract

Across the brain, circuits with continuous attractor dynamics underpin the representation and storage in memory of continuous variables for motor control, navigation, and mental computations. The represented variables have various dimensions and topologies (lines, rings, euclidean planes), and the circuits exhibit continua of fixed points to store these variables, and the ability to use input velocity signals to update and maintain the representation of unobserved variables, effectively integrating the incoming velocity signal. Integration constitutes a general computational strategy that enables variable state estimation when direct observation of the variable is not possible, suggesting that it may play a critical role in other cognitive processes. While some neural network models for integration exist, a comprehensive theory for constructing neural circuits with a given topology and integration capabilities is lacking. Here, we present a theoretically-driven design framework, Manifold Attractor Direct Engineering (MADE), to automatically, analytically, and explicitly construct biologically plausible continuous attractor neural networks with diverse user-specified topologies. We show how these attractor networks can be endowed with accurate integration functionality through biologically realistic circuit mechanisms. MADE networks closely resemble biological circuits where the attractor mechanisms have been characterized. Additionally, MADE offers innovative and minimal circuit models for uncharacterized topologies, enabling a systematic approach to developing and testing mathematical theories related to cognition and computation in the brain.

## Introduction

The brains of species from insects to mammals contain circuits specialized to represent and integrate continuous variables (Figure 1A) [1, 2]: the head direction circuits in mammals [3, 4, 5], fish [6], and flies [7, 8, 9, 10], the oculomotor system of vertebrates [11, 12, 13, 14, 15], and grid cell networks in mammals [16, 17, 18] (see Figure 1B,C,D). These circuits receive velocity inputs, representing the rate of change of the represented variable, and update their internal state in proportion to the instantaneous velocity [1]. The oculomotor circuit integrates head velocity signals to counter-rotate the eyes and hold the gaze fixed during head movements [15, 11]; it also integrates saccadic velocity commands to generate stable fixations at different gaze angles between saccades [13]. In the head direction and grid cell circuits for spatial navigation, self-movement cues from turning and walking update the internal pose estimates [5, 19, 20, 21, 22]. This so-called path integration (PI) computation underpins behaviors that are core for survival [23, 24].

**Figure 1:**
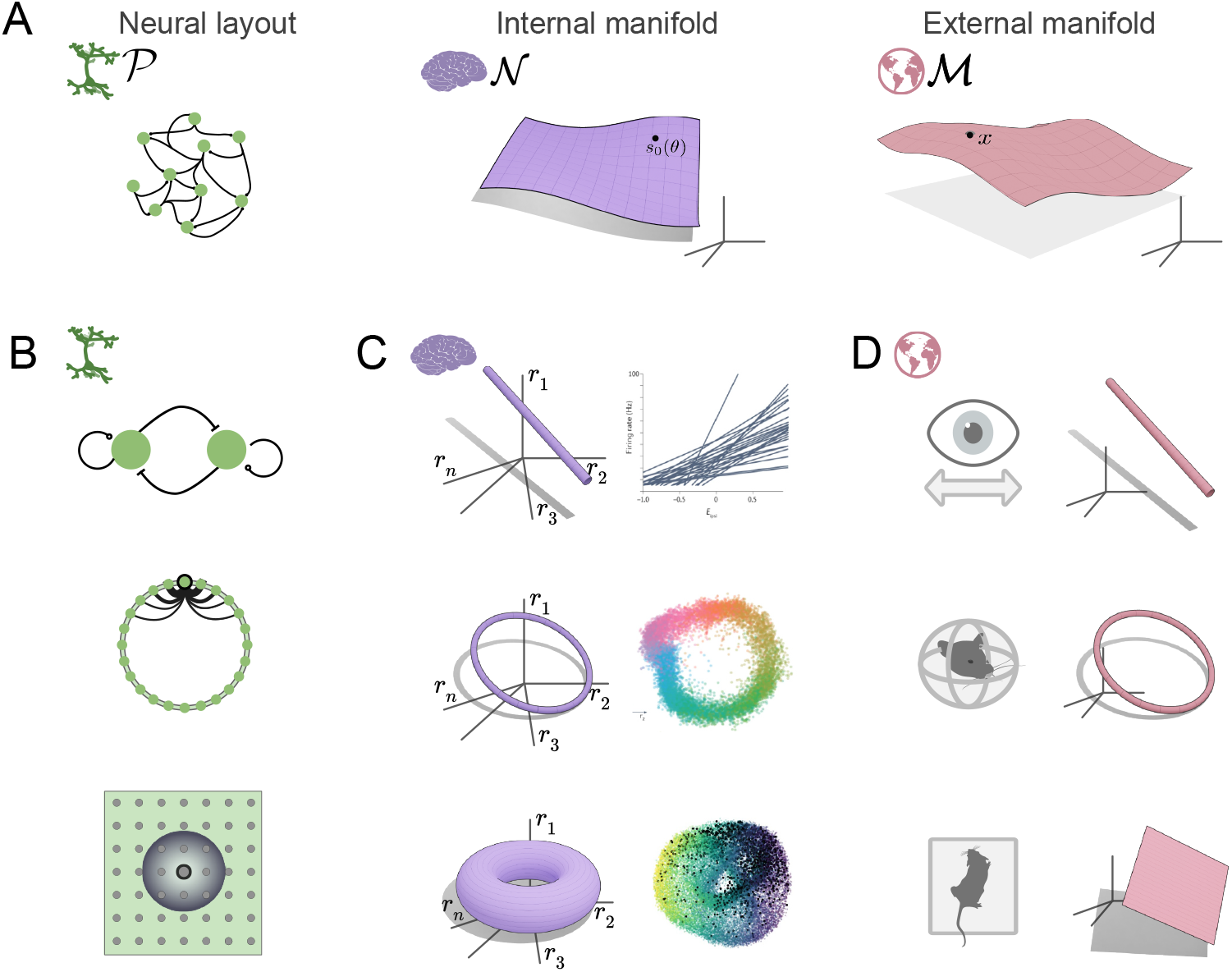
Existing continuous attractor network models and the biological systems where they are found. **A**. Left, schematic representation of a spatially embedded set of neurons and their connections. The neural connectivity constrains the patterns of neural co-activation, thus determining the dimensionality and topology of neural activity in the state space. Center, schematic representation of neural activity states, in this case forming a continuous manifold in state space. Right, schematic representation of the states of a (latent) variable in the external world. **B**,**C**,**D**. Examples of integrator circuits. Top row, integration in the oculomotor system. Center row, head direction system. Bottom row shows the grid cell system. **B**. Schematic representation of CAN models architecture for line, ring and torus attractors. **C**. Schematic illustration of the continuous manifolds of fixed points predicted and found to exist in the corresponding circuits, adapted from published work [15, 14, 34, 47, 35, 3, 48, 18, 46, 44]. **D**. Schematic illustration of variable manifolds.

Integration may also underlie the representation and mapping of other continuous domains including auditory sound spaces, parametric image variations, and emotional/aggression states [25, 26, 27, 28, 29, 30, 31, 32], and thus support inference, reasoning, planning, and imagination in all these domains.

Neural network models of these integrator circuit fall under the category of continuous attractor networks (CANs) [33, 1, 34, 35, 36, 15, 18]. All continuous attractor models posit recurrent circuitry to generate a continuous set of states that persist in the absence of external inputs (continuous attractors). However, not all CAN models are integrators: integrators must additionally contain a mechanism for updating the internal state based on velocity inputs. CAN models generate extensive predictions about circuit connectivity, activity, population dynamics, and lesions, and have stimulated extensive experimental work across species and circuits to test their predictions. Core novel predictions of these models have subsequently been validated via physiology, imaging, and connectomics: the dynamics and connectivity of the oculomotor integrator [37, 14, 12, 38] have been shown to match the hypothesized circuit model in considerable detail. The one-dimensional ring attractor dynamics, including fixed point dynamics, isometric representation in the head direction circuit in mammals matches [3] the predicted population dynamics of the ring integrator models in detail [39, 35, 34]. In insects, the connectivity and physical layout of the head direction circuit form an actual ring [7, 10] and exhibit some of the shift-like asymmetries hypothesized by a subset of the ring attractor models [39, 35]. In the grid cell system, the invariant two-dimensional population dynamics [40, 41, 42, 43] and its localization [44] to the predicted torus of fixed point states [18, 45, 46] has been directly observed in experiments. Thus, when available, circuit models have propelled a conceptual understanding of the structure and function of the mechanisms involved in integration, memory, and control of continuous variables, and driven experiments that have confirmed their mechanistic hypotheses

These models have been hand-crafted through intuition and insight, individually for each circuit or system in the brain. It is remarkable that the corresponding biological circuits have been found possess a structure, in the population dynamics and when direct physical comparisons have been possible in the circuit architecture, that closely matches these models [37, 14, 12, 40, 41, 42, 43, 7, 10, 38, 1]. This suggests that mathematically guided and conceptually minimal models are well-matched to the biology of the brain. Yet we lack a general mathematical theory to allow researchers to automatically construct such models for other continuous variables of a given dimension and topology, to generate predictions for future experiments and for potential use in machine learning applications involving such input variables.

Recent efforts to overcome this limitation center on training networks via gradient learning to perform continuous integration tasks on the desired variable [49, 50, 51, 52]. However, the difficulties of this approach for the formation of continuous attractor networks is that it is inefficient, and the results are not usually interpretable. Specifically, training on *M* -dimensional manifolds requires of order *k*^ℳ^ samples [53, 54], scaling exponentially with manifold dimension. In the few cases where the results become interpretable, it is only through mapping onto the original “hand designed” models. The combination of these factors and the striking match between biology and the minimal hand-crafted models suggests that a set of simple and general mathematical principles are used by biology to build such circuits and if discovered, can be used to directly construct circuit models for integration of arbitrary continuous variables.

Here, we present such a small set of mathematical principles to directly construct minimal, interpretable continuous attractor networks that integrate variables of rich topologies and geometries. The theoretical framework converts directly into a practical recipe for constructing integrating attractor networks of desired dimension and topology. Existing integration networks known from biology appear as special cases of this framework. We name the method MADE (Manifold Attractor Direct Engineering). Thus, MADE can serve as a generally useful framework for making circuit predictions about connectivity and function in neural integrators not yet discovered, including in high-level areas that perform various cognitive tasks.

## Results

Integration is the task of estimating the value of some (potentially latent) continuous variable *x*(*t*), based on an initial condition and inputs conveying information about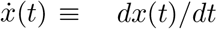, its instantaneous rate of change. For a variable to be integrable, it must be continuous and lie on a ‘differentiable manifold’: a smooth, continuous space that at small scales is similar to Euclidean space, though globally it may be non-Euclidean, with complex topology. For a neural circuit to integrate, its representations must form a differential manifold, and if the velocity signal is zero then the read out state should not change over time. In constructing a neural circuit that can integrate a given variable, we therefore need two components: a network that possesses a manifold of states that support a stable readout value, whose dimension and topology matches the variable, and a mechanism to allow velocity inputs to move states along the manifold. In what follows, we derive a general theory for achieving both with neural circuits, assuming that the stable readouts are stable population states on the manifold.

### Theory: Continuous attractor manifolds of desired dimension and topology

Here we describe the theoretical elements sufficient to construct a neural network possessing a continuous set of attractor states with desired intrinsic dimensionality *d* (e.g., *d* = 1 for a ring lattice and *d* = 2 for a plane) and desired topology specified by a manifold 𝒫.

Consider a set of *N* neurons and spatially embed them, equally spaced (in a lattice), according to the desired manifold topology 𝒫. With this embedding, each neuron has a unique *d*-dimensional coordinate *θ*_*i*_. This spatial organization is used for the specification of network connectivity, *W*_*ij*_ = *W* (*θ*_*i*_, *θ*_*j*_); it may but need not mirror the actual locations of neurons in neural tissue [18]. We use rate-based neurons with standard recurrent weighted sums and point-wise neural nonlinearity given by the function *f*. The activation of the neuron at *θ*_*i*_ is denoted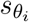. For better analytical transparency — so that weights and activations can be written as functions instead of lists of numbers — we follow others [34, 55] and take the continuum neural field limit. The discrete lattice of positions on the neural manifold 𝒫 and neural activations become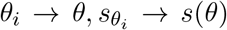, respectively. Additionally, ∑_*i*_ → ∫ *dθ*, ∑_*j*_ *W*_*ij*_*s*_*j*_ → ∫ *W (θ,θ*^′^)*s (θ*^′^)*dθ*^′^, so that the neural network equations are:

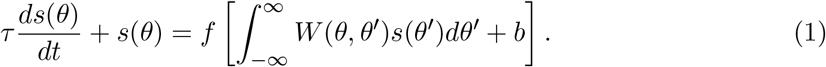

We will use the rectifying nonlinearity, *f* (*x*) = *x* if *x >* 0 and *f* (*x*) = 0 if *x* ≤ 0. Derivations that follow are conceptually and qualitatively independent of this continuum limit.

We seek interaction weights consistent with the formation, through symmetry breaking, of a single activity bump state that can be positioned anywhere on the neural manifold 𝒫. The set of such bump states will form the continuous set of attractor states of desired dimension and topology.

Let *W* be a kernel function, *W* (*θ, θ*^′^) = *k*(*d*(*θ, θ*^′^)), where *d*(*θ, θ*^′^) is a distance metric defined on 𝒫, and *k* is a continuous scalar function that is symmetric about the origin (see Figure 2A). Analogous to prior work [34, 48, 56], we set *k* to be locally excitatory and globally inhibitory. To avoid runaway excitability, we make it strictly inhibition-dominated (*k*(*d*) ≤ 0 for all *d*) as in [57, 18]; network activity can be non-zero because of a compensatory spatially- and temporally-constant excitatory feed-forward drive *b >* 0. Specifically, *k*(*d*) = −*k*_0_ + *k*_1_(*d*), where *k*_0_ *>* 0 is a positive number and *k*_1_(*d*) → 0 as *d* → *∞* with *k*_1_(0) = *k*_0_.

**Figure 2:**
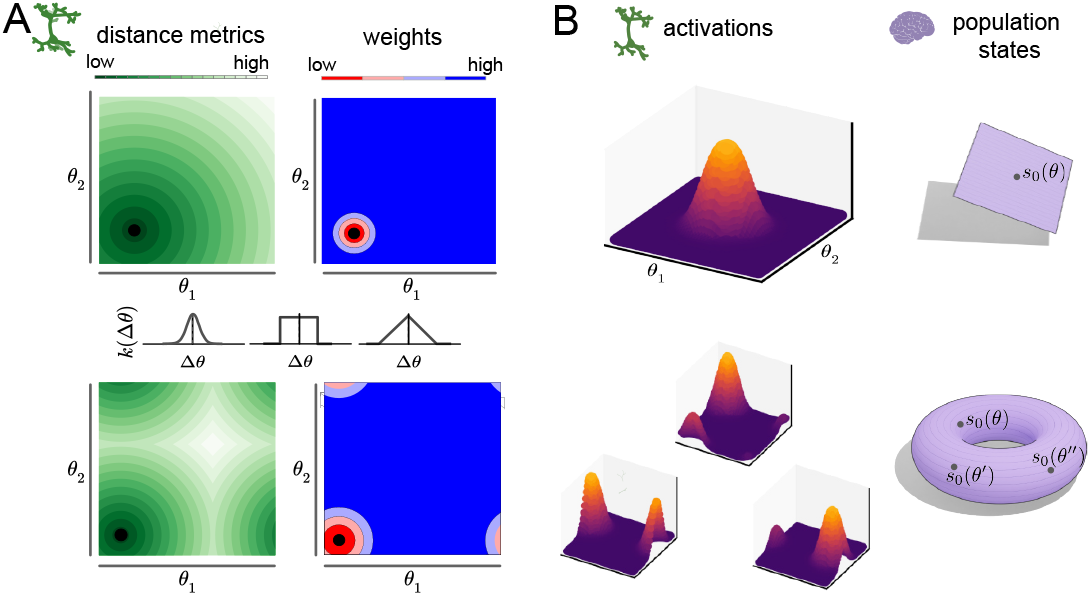
CAN construction and activity manifolds. **A**.Left, neural lattice 𝒫 for the Plane (top) and Torus (bottom) attractor networks. Black circles indicate the location of an example neuron, shades of green represent distance from other points on the lattice. Bottom right, inhibitory connectivity strength between the example neurons and all other points on the neural lattice. Middle inset, three examples of valid connectivity kernel functions k. **B**. Neural manifold in state space (top,right) and activity patterns on the neural lattice 𝒫 (top,left). Bottom row shows three activity patterns with bumps at different locations corresponding to different points on the activity manifold 𝒩.

Let the kernel’s length scale be given by *σ*, i.e., *k*_1_(*d*) *≈* 0 for *d* ≥ *σ*, with *σ* selected to be much smaller than the distances *L* over which the manifold 𝒫 has curvature. Thus, within any ball *V*_l_ of radius *l* such that *σ ≪ l ≪ L*, 𝒫 is flat. Since *σ* is the only spatial scale being introduced in the dynamics, we qualitatively expect that a localized bump state within the ball will have a spatial scale of O(*σ*). The conditions for the formation of a stable single bump state are thus the same as those for a globally flat manifold.

Since *W* is symmetric, Eq. 1 can be described through an energy function [58], and a stable steady state must exist. If the homogeneous state (all neurons equally active) were unstable, there must exist some other stable state, with broken symmetry. If the symmetry broken state is localized, we would refer to it as a bump state. Thus, we seek conditions under which the homogeneous steady state is unstable. The homogeneous steady state *s*(*x*) = *s*_0_ must satisfy

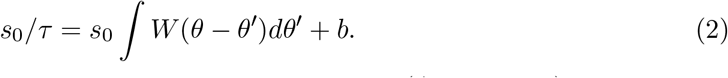

We derive the existence and stability of the homogeneous state (Appendix 1) following the analysis in Ref. [59], to obtain two requirements for the formation of a stable bump state:first, the Fourier transform of the kernel *k*_1_(*d*), which we denote as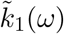, must be maximized at *ω* = 0; and second, this maximum must be larger than 1*/τ*. If *k* attains a positive maximum value at *ω* = 0, a rescaling can always make this maximum larger than 1*/τ*.

A broad sufficiency condition for the first requirement is if *k*_1_(*d*) ≥ 0 for all *d*, then its Fourier transform is maximized at zero (proof in Appendix 1). This condition does not include all interaction kernels *k*_1_ whose Fourier transforms are maximized at zero, but is a sufficiently broad class.

Thus, up to a rescaling of the strength of the interaction, an interaction *W* (*d*(*θ, θ*^′^)) will lead to the formation of a bump state if it can be rewritten as *W* (*d*(*θ, θ*^′^)) = *k*_1_(*d*(*θ, θ*^′^)) − *k*_0_ for: *k*_0_ ≥ 0; a kernel *k*_1_ that satisfies *k*_1_(*d*) ≥ 0 and *k*_1_(*d*) → 0 for *d* ≥ *σ* ; and sufficiently small *σ* over which the manifold 𝒫 is approximately flat. As a result, there is a set of stable fixed points corresponding to activity profiles that are in one-to-one correspondence with points on 𝒫: every stable single-bump activity pattern is centered at some point in 𝒫, and every point in 𝒫 forms the center of some stable single-bump state (see Figures 2B). Thus, the set of stable states of the dynamics in Eq. (1) form a continuous attractor manifold 𝒩 that has a bijection with the manifold of the neural layout 𝒫 and thus to the target manifold. Moreover, importantly for representation and integration of continuous variables, we show in Appendix 2 that 𝒫 and 𝒩 are isometric to each other, with respect to their intrinsic geodesic metrics.

### Theory: Integration on manifolds

The theoretical and practical frameworks outlined above show how to construct neural networks whose activity states possess a set of attractors forming a manifold 𝒩 of desired dimension and topology. Here, given the desired manifold 𝒩, we describe how the constructed attractor network with states matching the topology and dimension of 𝒩 can be augmented to endow them with the ability to perform velocity integration.

Note that to perform velocity integration of an external observed variable, the desired manifold 𝒩 may, but need not, coincide in dimension and topology with the manifold on which the observed variable states lie. This possibility is exemplified by grid cells, where the manifold 𝒩 of a grid module is 𝒩= 𝕋^2^ and is used to integrate animal velocities as animals move about in physical 2D space (thus ℳ= ℝ^2^). In a future work, we will consider the question of which internal manifolds 𝒩, not necessarily of the same topology or dimension as ℳ, permit accurate integration of velocities on ℳ. Here we show how to equip networks with attractor manifold 𝒩 with accurate path integration functionality for velocity inputs of matching dimensionality.

Previous models [34, 35, 18] constructed offset interactions between multiple copies of a continuous attractor network to permit external inputs to drive the state along the manifold. Here, we analytically derive the conditions required for an external input that has no knowledge about the structure and state of the continuous attractor network to generate appropriate movements along the nonlinear attractor manifolds of given topology, and show that offset interactions are necessary solutions.

For simplicity, consider a one-dimensional manifold with linear transfer function *f*. The stable bump states are fixed points of Eq. 1:

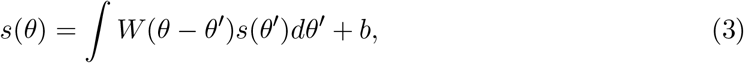

where *s*(*θ*) denotes an activity bump centered at any point in 𝒫. Consider two such activity bump states: *s*_0_(*θ*) centered at *θ*_0_ and *s*_0_(*θ* + *ϵ*) centered at *θ*_0_ − *ϵ*. For the neural state to move from *s*_0_(*θ*) to *s*_0_(*θ* + *ϵ*) in time Δ*t*, the time derivative *∂s/∂t* must equal

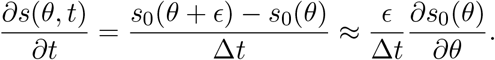

The movement speed is *v* = *ϵ/*Δ*t*. Multiplying by *τ* on both sides, we have

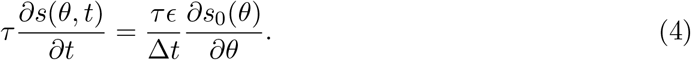

We can add 0 to the equation above, in the form (−*s*_0_ + ∫ *W* (*θ* − *θ*^′^)*s*_0_(*θ*^′^)*dθ*^′^ + *b*), which is zero because of the equality of Eq. 3), to obtain:

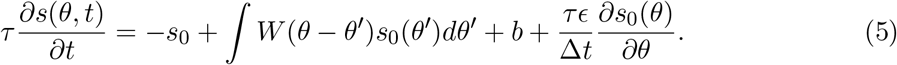

Comparing this expression to Eq. 1, we see that moving the bump with velocity *v* can be achieved by adding a feedforward input drive 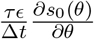 to the continuous attractor network. Though this appears to be a simple way to drive the activity bump on the manifold, it would require the external input to “know” the current value of 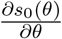, which varies along the manifold. Thus, the external input would need to know both the shape and current state on the internal neural activity manifold.

Observing that 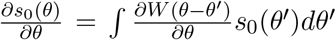 (from Eq. 3), and grouping like terms, we obtain

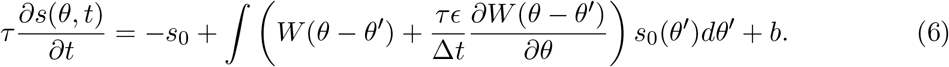

This expression has now “internalized” the desired input to move the bump, converting it into the weight asymmetry term 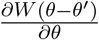, similar to [34]. The weight asymmetry is internal to the network, thus the velocity external input would not need to be aware of the internal state or shape on the attractor manifold to drive the bump. However, the external input would be required to dynamically modulate the degree of weight asymmetry, a biologically unrealistic requirement. As a final step, observe that for small *τϵ/*Δ*t* ≡ *δ*, by Taylor expansion,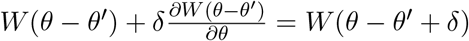 Thus, we obtain

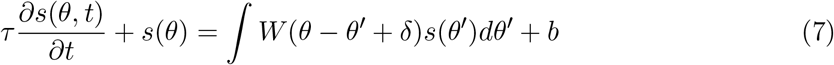

Because we have that *δ* = *τϵ/*Δ*t* = *τv*, the equation above results in a moving bump along the internal state-space manifold of fixed points 𝒩 with speed *v* = *δ/τ*, without any external velocity input or temporally varying modulation of network weights. The network corresponds to the original continuous attractor network constructed in the previous section, with the modification that the weights, instead of being symmetric, have a small offset in a particular direction *δ* along the neural circuit manifold 𝒫. The speed of bump movement on 𝒩 is proportional to the magnitude of the offset, |*δ*|, and inversely proportional to the neural time-constant.

This continuous-speed flow may form a periodic cycle on specific manifolds (e.g. Ref.[18, 35]). In these cases, the network is a limit cycle attractor. On generic manifolds, however, this flow need not close periodically on itself. The result will be a quasiperiodic attractor dynamics [60]. We therefore refer to these as Quasiperiodic Attractor Networks (QANs). The flow of activity patterns in a QAN defines a constant vector field Ψ on 𝒩.

For several attractor manifolds 𝒩 of dimension *d* (in particular, ‘parallelizable manifolds’ such as the Euclidean spaces ℝ^d^ and the Torii 𝕋^d^) it is possible to construct *d* QANs with linearly independent flows, and 2*d* QANs with two mutually opposing flows in each of *d* dimensions (defined by weight matrices *W* (*θ* − *θ*^′^ ± *δ*_m_), where *δ*_m_ is a displacement vector of norm |*δ*| along the *m*^th^ manifold dimension). Each sets up a constant vector field Ψ_±m_ on 𝒩. For these manifold topologies [34, 35, 18], opposing-pair QANs numbering 2*d*, where *d* is the manifold dimension, can generate smooth non-vanishing flows of any direction at every point and are thus sufficient to construct integrators. The combined dynamics is given by:

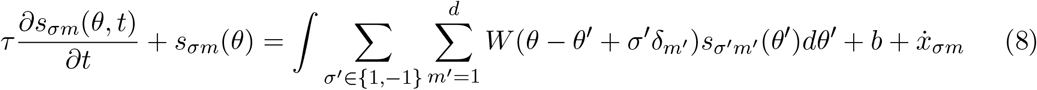

where *s*_σm_ indicates neural activities in the individual QANs and 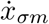 is an input carrying information about the rate of change of the external variable in the *m*^th^ direction.

Coupled in this way, the QANs form a network whose combined activity state moves on 𝒩 in a way controlled by the velocity inputs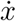, which modulate the activity levels of the individual QANs. When 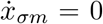 for all *σ, m*, the action of the opposing QANs along each dimension restores the symmetry of the system and *s* remains stationary (it does not flow along 𝒩). Otherwise, the terms 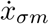 differentially modulate the activation of the QANs, causing the activity bump on 𝒫 to flow in the direction of the positively modulated QANs. The result is a time-varying vector field Ψ_*t*_. For accurate path integration, the component vector fields must be smooth and the set of QANs must generate a complete basis set of non-vanishing vector fields at every point on 𝒩. This condition is satisfied by using 2*d* QANs for Euclidean spaces ℝ^d^ and Torii 𝕋^d^, thus the prescription above is sufficient for integration on these manifolds.

On other manifolds, 2*d* opposing QANs for the *d* manifold dimensions are not sufficient for accurate integration. For instance, in the case of even-dimensional spheres, the hairy ball theorem states that every continuous tangent vector field must vanish at some point(s) [61, 62, 63]. In other words, a continuous vector field Ψ_±m_ generated by the QAN prescription above will be zero somewhere on the sphere; at that location, the QAN will not be able to drive bump movement; thus, *d* QAN pairs will not suffice for good integration everywhere. Further, on non-orientable manifolds such as the Möbius band, it is not possible to define continuous vector fields that are globally orthogonal everywhere and smooth. Thus, while the approach above provides a unified way to construct integrating continuous attractor networks — including all those with a single bump state currently found in the neuroscience literature [34, 35, 47, 48, 56] — it needs to be further generalized for manifolds that do not permit non-vanishing continuous tangent vector fields everywhere.

### Generalization: Killing vector fields

To enable accurate path integration over a significantly wider set of manifolds (excluding the Klein bottle), we now broaden and further generalize the concepts developed above. The approach replaces the constant weight offset vector fields Ψ_±m_ with the more generally applicable Killing vector fields [62]: Killing fields are vector fields on a manifold whose flows preserve the structure of the manifold, i.e., they are continuous isometries on the manifold. Conceptually, if each point of an object on the manifold is displaced by the corresponding Killing vector, it will move without distortion. Killing fields form a ‘vector space’, such that linear combinations of Killing fields are also Killing fields. The manifold isometric property of Killing fields means that activity patterns are rigidly translated over 𝒫 through the flow Ψ_*t*_ without changes in area, a necessary condition for accurate integration [34].

To generate Killing fields in each QAN, the constant weight offsets are replaced by an appropriate position-dependent offset:

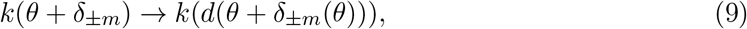

where ±*δ*_m_(*θ*) is the offset vector of the *σ, m*^th^ QAN at coordinates *θ* on 𝒫. This allows for weight offsets to vary at different locations on the manifold 𝒩 consistent with non-constant

Killing fields required on the sphere (Figure 3C). This simple change, and allowing the number of QANs to be larger than 2*d*, endows a much broader class of continuous attractor manifolds including spheres and Möbius band with integration functionality. For a two-dimensional sphere, three basis Killing fields (*d*_*kill*_ = 3) are required (each corresponding to rotational symmetry along one principal axis; Figure 3C). Although each field vanishes at two points on the sphere, at least two fields are non-vanishing and point in independent directions along the manifold at any point, forming an overcomplete basis such that it is possible for the network to perform accurate path integration.

**Figure 3:**
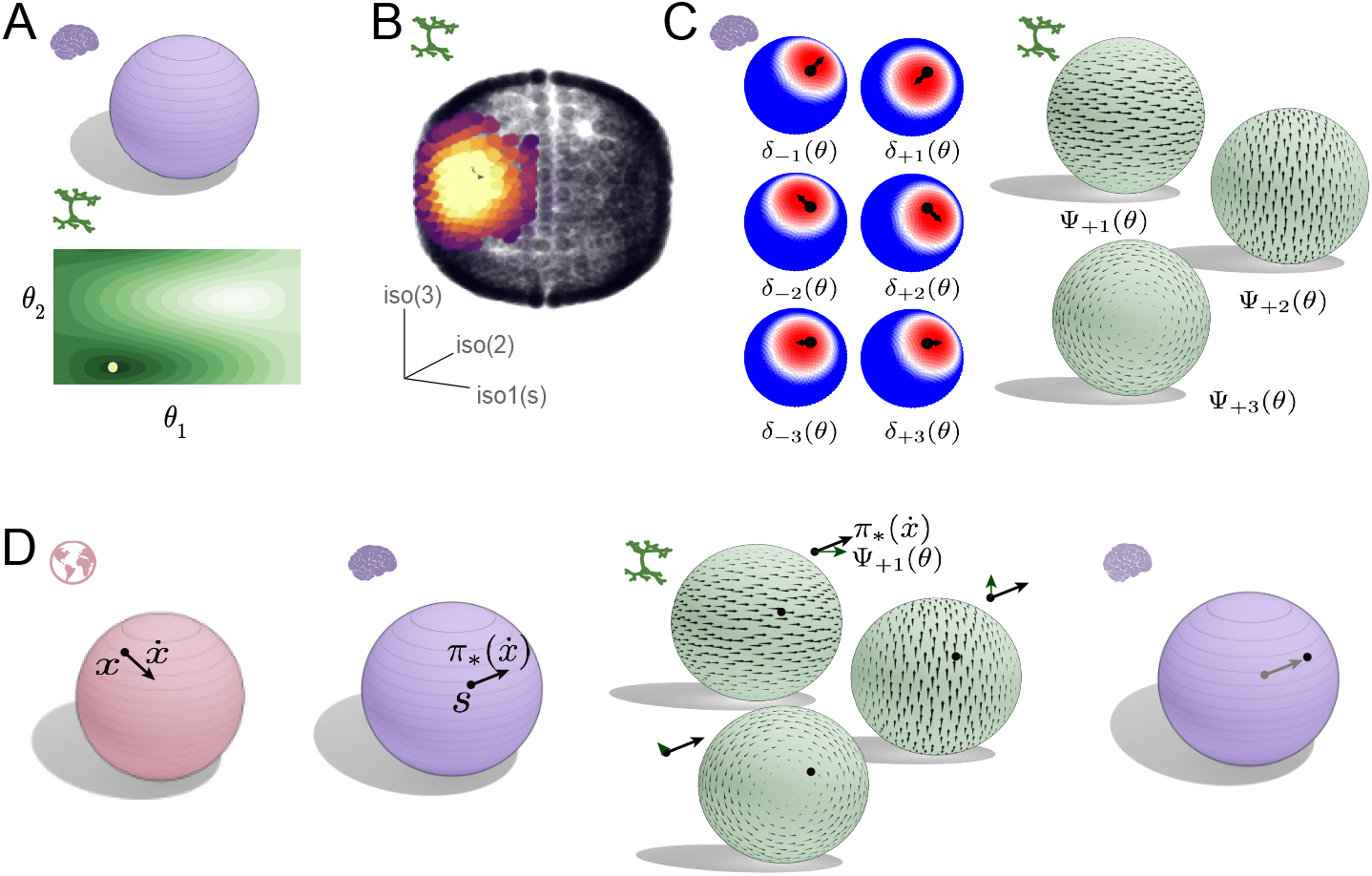
Quasiperiodic Attractor Networks for Path Integration. **A**. Schematic representation of a desired 2D spherical set of fixed points in state space and corresponding connectivity on 𝒫. **B**. Example activity bump plotted on the neural manifold 𝒫. **C**. Schematic illustration of Killing vector fields for the sphere manifold, left, and resulting offset connectivity weights on 𝒫, right. **D**. Schematic illustration of the QAN approach to velocity integration. Left two panels, relationship between changes in the variable on ℳ and on the neural 𝒩 manifold, and associated tangent vectors. Center, each QAN receives a velocity-dependent input based on the tangent vectors at left projected onto its Killing fields, and the activity of all networks is combined. Right: this results in a trajectory in the state-space 𝒩, which corresponds to velocity integration of inputs from ℳ.

Finally, we generalize how an external manifold ℳ may be mapped to the internal integrating manifold 𝒩, by mapping velocity vectors in the external space to the QANs within the network. Throughout, our construction seeks to make 𝒫 and 𝒩 isometric, and indeed they are, as shown in Appendix 2. However, as noted at the start of this section, 𝒩 need not exactly match the topology of the external variable: 𝒩 = T^2^ of a grid module represents positions on ℳ = ℝ^2^ of the externals partial variable. Similarly, the dimensionality of 𝒩 could equal or exceed that of ℳ: a planar integrator network is capable of integrating an external one-dimensional variable if the velocity inputs are one-dimensional. For instance, grid cell responses on a linear track appear to be generated as a slice through their 2D manifold of states [41, 64].

Define *π* as the mapping of ℳ to 𝒩 (which can be the identity map or the isomorphism map when ℳ and 𝒩 are isomorphic, such as when head direction is represented in a ring attractor, or a many-to-one map as when spatial position is represented in a single grid module). The Jacobian *π*_⋆_ is a map from the tangent space of ℳ to the tangent space of 𝒩: it is the operator that maps tangent vectors from ℳ (i.e.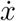) to tangent vectors of 𝒩 [63, 65]. In other words, the velocity vector 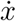 is ‘pushed forward’ through the map *π* into 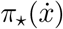 (Figure 3D). The coupled system dynamics can be written as

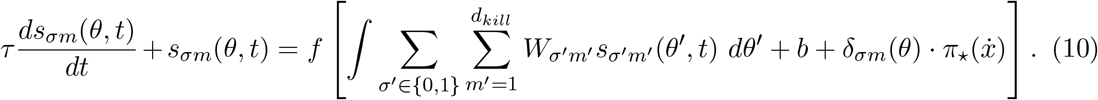

where *W*_σm_ refers to the Killing-field weights from Eq. 9, *d*_k*i*ll_ defines the minimal number of independent Killing fields. The term 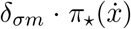 refers to the projection of the velocity pushed through *π* onto the (*σ, m*)^th^ QAN. Note that in general, the Jacobian *π*_⋆_ maps the tangent space at a specific point on ℳ to the tangent space at a specific point on 𝒩, making it dependent i principle on both *x* ∈ ℳ and *s* ∈ 𝒩. Thus, neural circuits generating 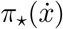 would require access to both the integrator’s neural state and the external variable. While the neural state *s* is available to the brain, *x* cannot be directly observed. However, if the integrator network maintains an accurate estimate of this variable — an expected property of a reliable integrator — then the brain can instead evaluate *π*_⋆_ at the integrator’s state on ℳ as a proxy for *x*.

The constant vector fields on the ring and torus manifolds described above (and effectively discovered in previous work) are Killing fields. Therefore, this approach encompasses previous work and provides a broader general framework for constructing minimal biologically plausible continuous attractor neural networks capable of path integration on spaces of various dimension and topology. Next, we demonstrate how to practically construct the networks, the examine the effectiveness of the approach through extensive numerical simulations of path integration in MADE integrator networks.

### Practical construction of CAN integrators with MADE

With the complete conceptual and mathematical frameworks in place, we now illustrate through numerical simulation how to apply the MADE prescription to construct various CANs and integrators of desired dimension and topology. The simulations also allow us to validate the functionality of the resulting CANs and integrators. For simplicity, here we focus our description on one and two-dimensional surfaces, allowing us to construct line, ring, plane, cylinder, torus, sphere, Möbius band and Klein bottle topologies and geometries (Figure 4A). The procedures outlined here can be straightforwardly generalized to apply to manifolds of different dimensionality.

**Figure 4:**
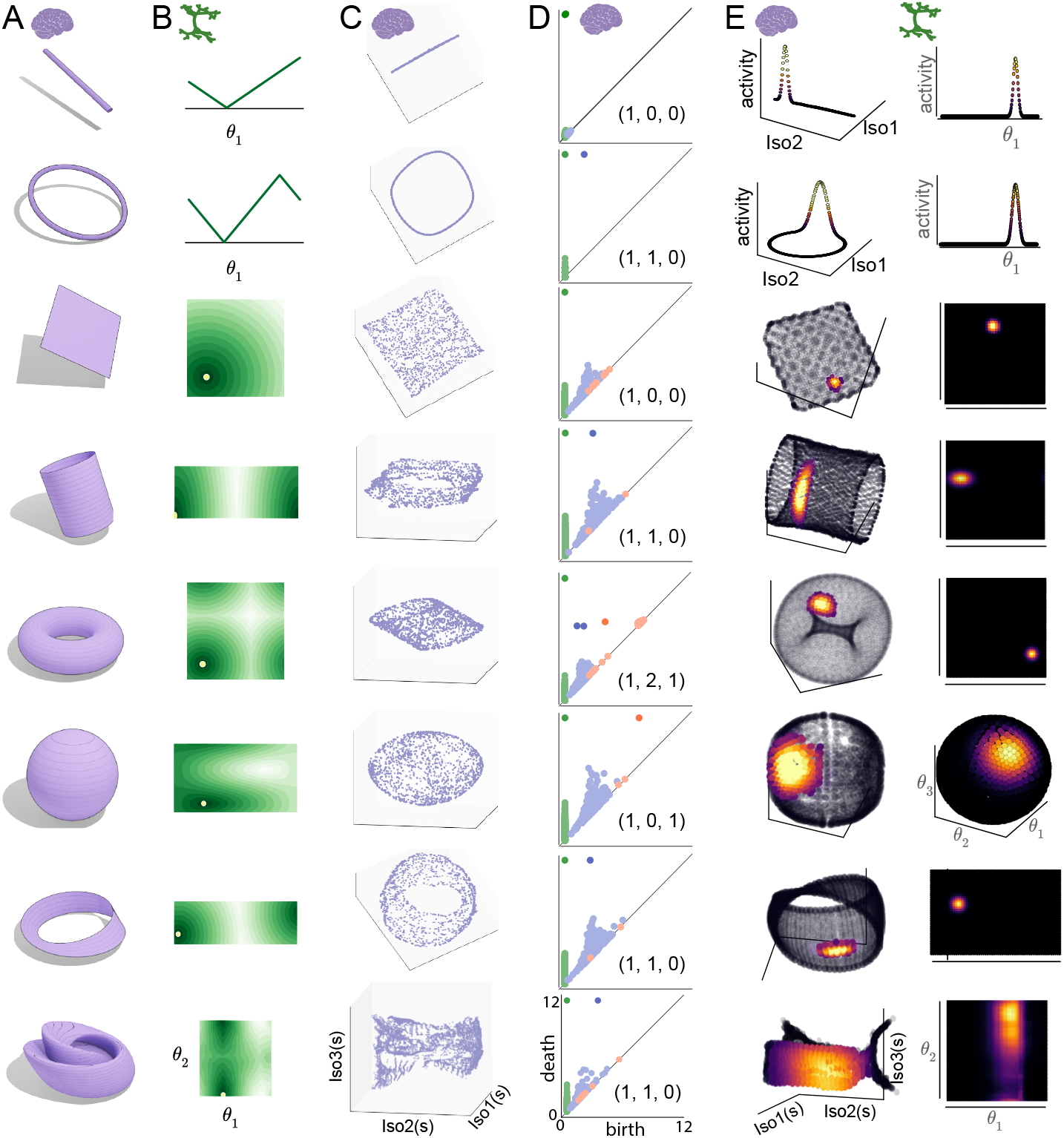
Stationary states and manifold topologies of the MADE CANs. **A**. Desired population activity manifold topology for CANs constructed with MADE for several manifolds (from top to bottom): line, ring, plane, cylinder, torus, sphere, Mobius band and Klein bottle. **B**. Distance functions over the neural lattice 𝒫 for selected example neurons. **C**. Low dimensional embedding of the neural activity manifold 𝒩. **D**. Betti number and persistent homology bar code for each CAN’s neural population states (in 𝒩). **E**. Left: Activity of one example neuron over 𝒩 (low dimensional embedding). Right: Stationary population activity states form localized bumps on the neural lattice 𝒫.

We first construct a neural surface 𝒫 that is isometric to the target state-space manifold 𝒩. For the sphere attractor, we construct 𝒫 as an embedding of the two-dimensional unit sphere in ℝ^3^, and for the Klein bottle attractor 𝒫 was an embedding of a finite cylinder manifold with appropriate identification of the cylinder end-points to each other in ℝ^4^). For several other manifolds (including all others from Fig. 4), which admit a flat metric, we define a rectangular two-dimensional space [0, *L*_1_] × [0, *L*_2_] (Figure 4B) and provide an appropriate distance function on the rectangular space. For example, for the torus manifold, *L*_1_ = *L*_2_ = 2*π*, and distances are computed respecting the periodic boundary conditions that identify 0 and 2*π* as the same point.

Given 𝒫, we next approximately evenly place neurons on the surface. For manifolds with a flat metric, this involved placing neurons on an *n* × *n* rectangular lattice on this space, where *n*^2^ is the total number of neurons. For the sphere, we spaced neurons at regular intervals along a Fibonacci spiral over the unit sphere (see Methods)to approximate an even placement on the sphere. Thus, for each neuron we define their 𝒫 coordinates *θ*_*i*_.

Next, we computed the connectivity of the network *W*_*ij*_, which depends on the (geodesic) distances *d*(*θ*_*i*_, *θ*_*j*_) between pairs of neurons with coordinates *θ*_*i*_ and *θ*_*j*_ on 𝒫. With appropriate coordinate parametrization for the neurons, these geodesic distances can be computed via analytical expressions (for instance, as Euclidean distance with periodic boundary conditions on a torus attractor), or via a simple numerical computation (see Methods). Connectivity is then given by the *n*^2^ × *n*^2^ matrix with entries *W*_*i*,j_ = *k*(*d*(*θ*_*i*_, *θ*_*j*_)), where *k* is a kernel function (Figures 2A, 3A, 4B) satisfying the requirements described earlier for the formation of activity bump states (see also Appendix 1). We used a scaled Gaussian kernel such that the connectivity between pairs of neurons was strictly negative and *W*_*ij*_ = 0 if *d*(*θ*_*i*_, *θ*_*j*_) = 0 (see Methods). Other choices of kernels yield similar results (data not shown). Neural activity is simulated based on these weights according to Eq. 2. We will provide Python and Julia code that implements the MADE prescription for CANs (see Methods).

### Validation of CAN states and dynamics

To validate the MADE CANs, we first characterize where the states of the constructed networks localize. To do so, we sample population activity data from each model by randomly initializing each network and allowing the initial state to settle to a stationary state (see Methods). This state forms one population vector sample; we repeat the process 2500 times for each network. We apply nonlinear dimensionality reduction via ISOMAP, which has proven useful for the visualization of nonlinear low-dimensional manifolds in real data [3], to the resulting point cloud of stationary population activity states. The resulting structures (Figure 4C) visually matched the desired manifolds (Figure 4A): the population responses of the MADE CANs localize to low-dimensional sets of states that appear homeomorphic to 𝒩.

To quantify the structure of the resulting population states, we use persistent homology, a Topological Data Analysis [66, 67] technique that has been applied with success in neuroscience [68, 3, 44]. Persistent homology supplies Betti numbers that characterize the topology of the set of stationary states of each network (see Methods). Betti numbers catalog the number of “cavities” of each dimension present on a manifold; the first three Betti numbers correspond to the number of connected components, rings and two dimensional cavities, respectively. Betti numbers don’t provide a complete or unique description of manifold structure (e.g., the ring and the cylinder share the same Betti numbers while having different dimensionality), but they provide a quantitative confirmation that the MADE CANs match their intended targets. The Betti numbers of all MADE CANs population states match those of their target manifolds (Figure 4 D).

We next visualize the instantaneous population activity states as functions on the neural lattice. The localized kernel connectivity on the manifold was expected to stabilize single activity bump states on the manifold. A stationary population activity state can be directly visualized on the neural lattice by coloring neurons according to their activity level. Indeed, we see that the stationary population states correspond to localized bumps of activation on the neural lattice 𝒫 and activity manifold 𝒩 (Figure 4E).

Next, we characterize the intrinsic dimensionality [3, 69] of the stationary states of the MADE CANS. Intrinsic dimensionality at a point on a manifold is the numbers of degrees of freedom of movement along the manifold at that point. Intrinsic dimensionality would allow one to distinguish, for example, a ring (one dimensional) from a cylinder (two dimensional). Dimensionality is generally a difficult (and ill-posed) quantity to estimate in noisy data, and existing works use various methods [3, 70, 71, 72, 73]. For MADE CANs, which we can run in a noiseless setting, intrinsic manifold dimension is well-defined.

We adopt an approach [73, 72] based on estimating the dimensionality of the tangent space to a manifold (see Methods)(Figure 5 A, left). The tangent space *T*_*s*_ 𝒩 at a point *s* ∈ 𝒩 is the best linear approximation of the manifold at that point and has the same dimensionality as the underlying manifold [74, 63, 65]. We consider the set *S* of points in a small neighborhood of *s* (see Methods)and apply PCA to determine the number of large principal components needed to describe the data. This gives us the dimensionality of the tangent space at *s* and thus the local intrinsic dimension of the manifold.

**Figure 5:**
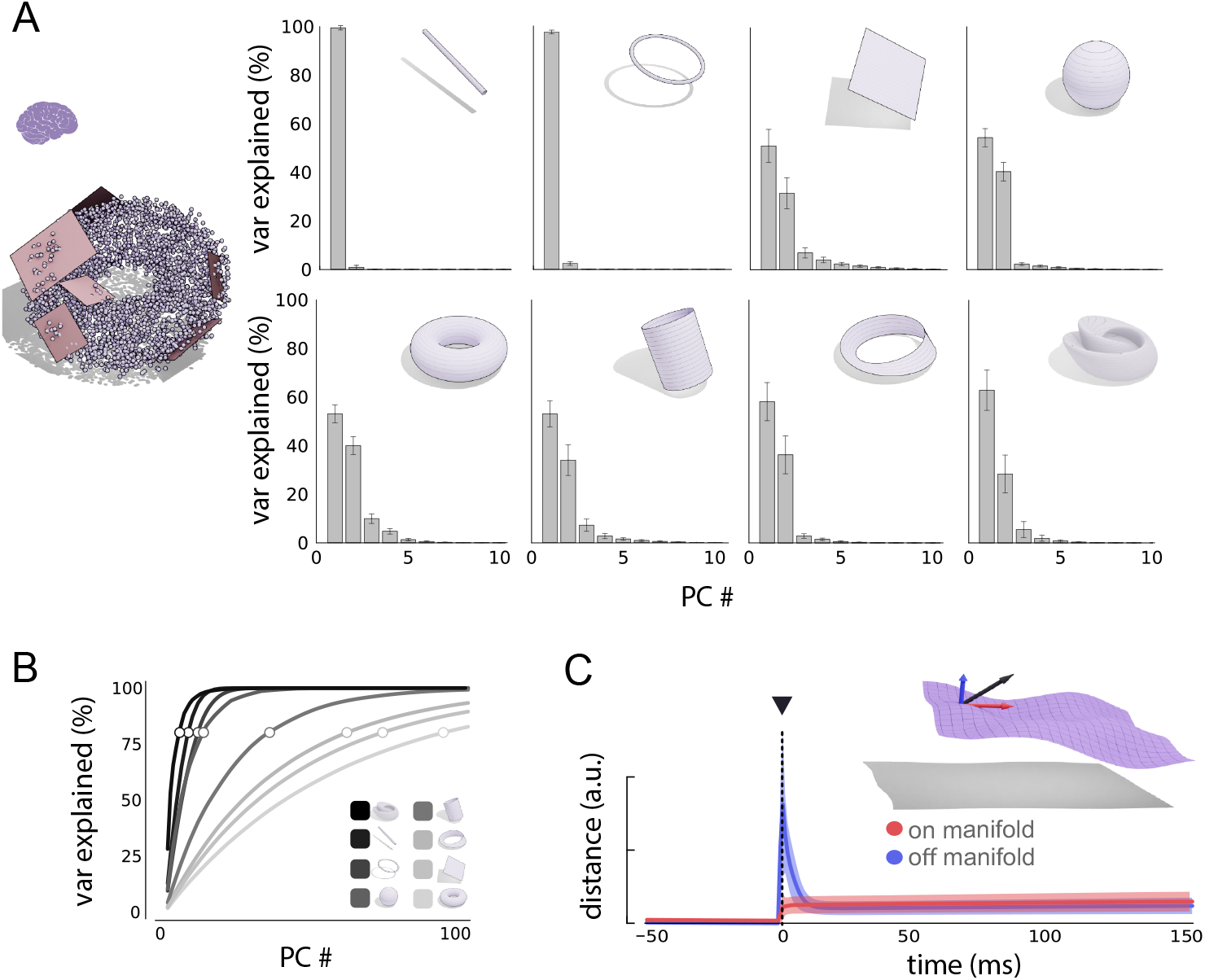
Dimensionality and attractor dynamics of the MADE CANs. **A**, Left, tangent planes approach to computing the intrinsic manifold dimension (schematic) of 𝒩. Right, estimated tangent space dimension for each manifold, which estimates the low intrinsic dimensionality of the CAN networks. **B** Cumulative manifold variance explained by global PCA analysis: the slow saturation of the curves shows that the linear (embedding) dimension of the manifolds can be large. **C** Numerical simulations to probe attractor dynamics. Inset: activity manifold, perturbation vector (black) and on-manifold (red) and off-manifold (blue) components of the perturbation. Main plot: Time-varying distance from the starting point in the off-manifold and along-manifold dimensions.

Repeating this analysis across multiple randomly selected sample points *s* for each MADE CAN, we confirmed that all manifolds had the expected intrinsic dimensionality given their topology: line 1 ± 0.0 (mean ± standard deviation, across multiple repeats), ring: 1 ± 0.0, torus: 2 ± 0.0, sphere: 2 ± 0.0, Möbius band: 1.96 ± 0.16, cylinder: 2 ± 0.0 and plane: 2.05 ± 0.23 (Figure 5 A). By contrast to the small intrinsic dimensionality of the constructed CAN manifolds, their extrinsic linear dimensionality, estimated by the minimum number of principal components required to represent the manifold as a whole, is large (Figure 5 B).

Finally, we examined whether the stationary manifolds of the MADE CANs are neutral attractor states, with rapid decay of off-manifold perturbations, together with no state drift along the manifold in the absence of noise and external inputs [34, 1]. First, we consider manifold stability by computing Betti numbers of the population states in networks simulated with varying noise conditions, and find that except in the most severe noise case, we recover the same Betti numbers for the noisy dynamics – indirectly showing that the manifold is attractive and robust to noise (see Methods)(Figure S1). Second, we more directly perturb the neural population state with a randomly oriented vector of fixed magnitude (see Methods), repeating this experiment for multiple initial states and random perturbations, and observe the dynamics by which the perturbed state evolves. To quantify on- and off-manifold dynamics following perturbation, we again used PCA to estimate the manifold’s tangent space in the neighborhood of the initial state. The distance between the perturbed and initial (pre-perturbation) states along the tangent space dimension was considered the on-manifold perturbation component; the rest (along the remaining *N* −*d* dimensions) was the off-manifold perturbation(see Methods). We find very limited on-manifold drift and strong decay of the off-manifold component of the perturbation, as intended (Figure 5 C).

### Practical construction of integrators with MADE

To generate the QANs that combine to create neural integrator circuits, we slightly modify the connectivity structure of MADE CANs. We start with the same procedure as before to construct 𝒫 and compute the distance function *d*. For a QAN indexed by *σ, m* we simply apply a shift 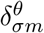 to the *i*^th^ neurons coordinates before computing *d* such that 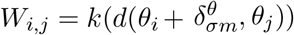. For some manifolds with a flat metric (e.g. plane, torus) 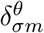 was identical for all points *θ* ∈ 𝒫 and was taken to be a vector of magnitude |*δ*| oriented along the *m*^th^ direction on 𝒫. In others (e.g. the sphere), the offset vector varied as a function of position along the manifold. For each dimension *m* we defined a Killing Vector field Ψ_±m_ and evaluated it at *θ*_*i*_ to obtain the offset vector (see Methods). Given an external velocity signal for a trajectory on ℳ, we use the map *π* from ℳ to 𝒩 to obtain the inputs to each QAN. Network activity is simulated based on weights and these inputs according to Eq. 10. We will provide Python and Julia code to implement the MADE prescription for neural integrators (see Methods).

### Validation of MADE integrators

To examine the performance of each MADE integrator in representing and tracking time-varying external variables, we provide the circuit with the velocity of a simulated random trajectories of the variable *x*(*t*) ∈ ℳ and track how the network’s internal state changes. We first consider how the firing of a single cell varies with the external variable, by plotting its tuning curve or firing response as a function of the external variable, estimated over a long velocity trjectory (Figure 6A). The existence of a localized activity bump (Figure 6A, top three panels) means that the circuit has correctly inferred external position: the cell fires at a specific position and not other random positions, and the network has transferred the internal bump activity pattern into a corresponding pattern as a function of location on the external manifold ℳ. In cases where the external manifold ℳ is not isomorphic to the internal manifold 𝒩, such as when a plane in ℳ is represented by a cylinder or a torus in 𝒩, a continued linear trajectory along one direction in ℳ corresponds to a periodic traversal on 𝒩, and thus one would expect repeating bumps in the tuning curve along that dimension, as we find (Figure 6A, panels 4-5). Note that based on the details of how we periodically connected the boundaries of our rectangular neural lattice to obtain a torus, we would obtain a square grid tuning curve (as shown) or a triangular grid tuning curve (as previously described for grid cells in [48, 56]). Finally, the tuning curves for the sphere and Mobius strip are single bumps, as expected (Figure 6A, last two panels).

**Figure 6:**
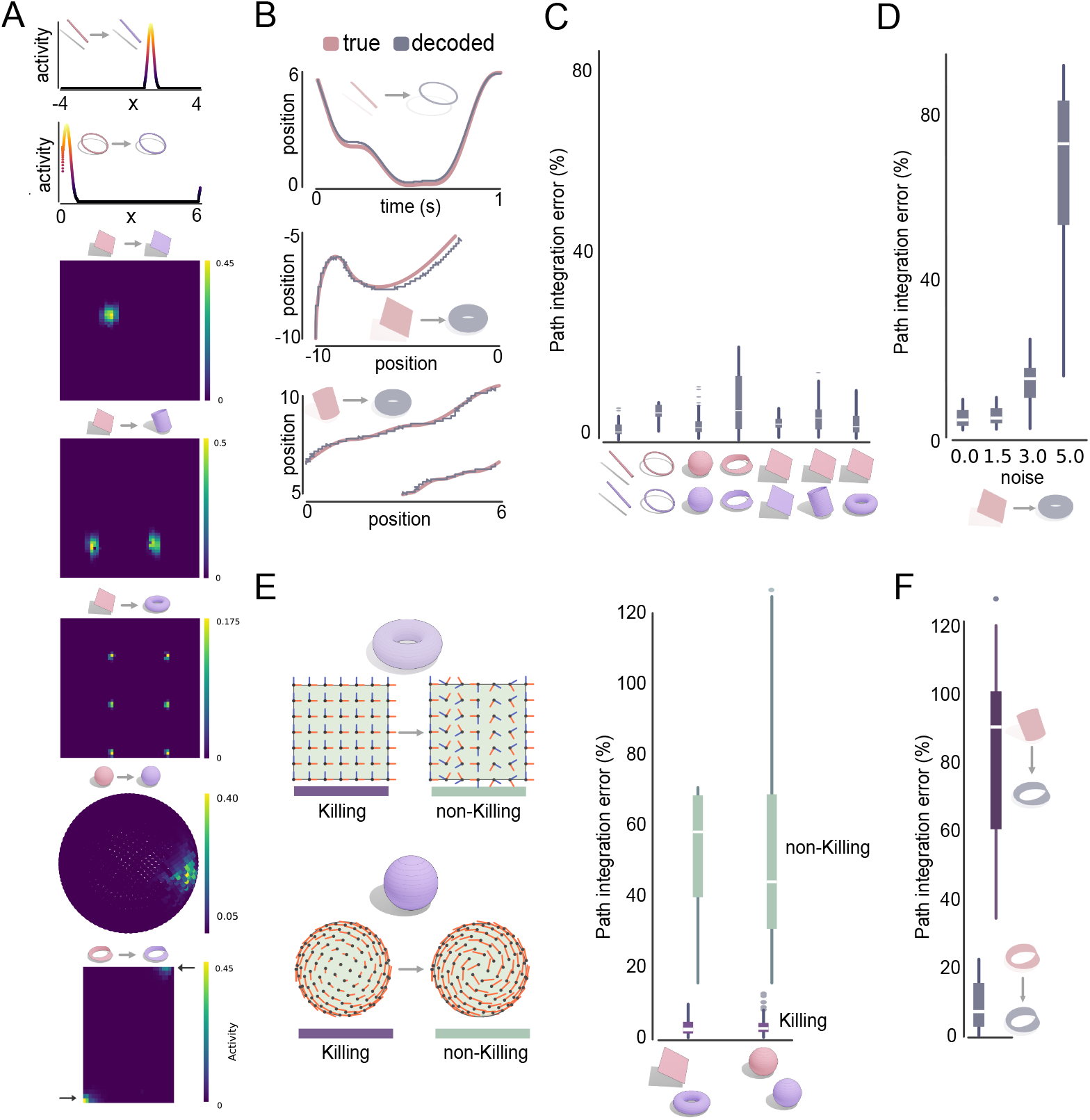
Numerical simulations of path integration performance with MADE path integrators. **A**. Tuning curves of single example neurons as a function of the external (latent) variable x. Insets show the manifold topologies of the external variable (red) and neural population states (blue): these pairings might be of identical manifolds, or e.g. a 2D Euclidean manifold in x could be mapped to a cylinder or torus, etc. in the neural population states. **B**. Example input trajectory (red) and decoded trajectory from the neural population response (blue). **C**. Decoding error across multiple simulations for various external-neural manifold pairs. Decoding error is shown as percentage of trajectory length over ℳ. Colored boxes show the interquartile range, white lines the mean, circles outliers and vertical lines the 95^*th*^ percentile confidence interval. **D**. Same as **B** but for torus attractors with varying amounts of noise. **E**. Left: Killing and non-Killing weight offsets for the torus (top) and sphere (bottom). Right: Same as **C** for integrators correctly constructed with Killing weight offsets, and with the non-Killing weight offsets from the left. **F**. Same as **C** for Möbius to Möbius (left) and cylinder to Möbius mappings (right).

We can more directly quantify how closely the network tracks the external variable *x*(*t*) by decoding it from the network’s internal state **s**(*t*), as 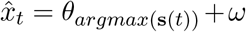 where *ω* is an offset used to account for the fact that, in some cases, 𝒩 was periodic while ℳ was not (e.g. torus and plane, respectively) (see Methods). When ℳ and 𝒩 are chosen such that *π* is either an identity map or a periodic mapping, the networks show very accurate integration over periods of several seconds of simulated activity (Figure 6B,C). Decoding error remains low even in the presence of moderate noise (Figure 6D) (see Methods). Thus, MADE networks support accurate integration, even in non-trivial scenarios such as the cylinder-torus manifold pairing and even on the Möbius band manifold, which have not been described previously.

We performed additional experiments on circuits requiring Killing vector fields to integrate. To show the necessity of Killing fields, we built torus (𝒩 = torus, ℳ = plane) and sphere (ℳ = 𝒩 = sphere) integrator networks, but varied the QAN weight offsets relative to the Killing field prescription. For the torus, we varied the orientation of the offset vectors, while for the sphere we changed their lengths to be of constant magnitude everywhere (except at two poles, where the magnitudes were left at 0), (see Figure 6E, left), (see Methods). The constant-magnitude non-Killing field on the sphere may be considered a direct extension of the constant offset vector fields used for flat manifolds and used in all prior work in the construction of neural integrators. In both cases, we observed a dramatic deterioration in integration accuracy, Figure6E (right). The result underscores the importance of Killing fields for integration on manifolds with a non-flat metric.

Finally, we considered integrating velocities from a cylindrical external variable on a network with Möbius band topology. Both manifolds are two-dimensional with one periodic and one non-periodic dimension. However, while a rectangle is glued without a twist to make a cylinder (which has two surfaces, inner and outer), it is glued with a twist to make a Möbius band (which has a single surface) with the consequence that there is no simple continuous mapping between the two. Proceeding naively by simply mapping the two manifolds onto each other by ignoring the flipped boundary of the Möbius band, it is unsurprising that integration is significantly less accurate (Figure6F). In future work, it will be interesting to consider which pairings of external to neural manifolds will provably permit accurate path integration.

## Discussion

### Summary

Here, we have presented MADE, a mathematical theory and recipe for constructing biologically plausible neural CANs and integrator networks of desired topologies and geometries, with single- or multi-bump tuning curves. The mathematical theory unifies existing biologically plausible continuous attractor and integrator models involving bump-like activation functions, which emerge as specific cases of the MADE theory.

The theory provides a first-principles derivation showing that multiple copies of a basic network must be coupled together for integration with biological constraints, in part to relieve demands for rapid synaptic modulation and in part to remove the need for velocity estimating regions from knowing the full nonlinear structure and current state of the integrator network. It also predicts that manifolds without a flat metric will require an overcomplete set of network pairs in the form of QAN networks, relative to the intrinsic dimensionality of the manifold: thus, integration on a two-dimensional spherical surface requires more than 2 QAN pairs.

We envision MADE to be useful to distinct fields: for deep-learning models that might require accurate low-dimensional neural network attractors and integrators, and for neuroscience, where MADE provides de novo models and novel circuit-level mechanistic predictions for the structure of other possible integrators in brain that may be uncovered in the future.

Indeed, given recent discoveries that path-integrating neural circuits generalizably represent multiple cognitive variables, it is likely that such circuits are used by the brain to perform cognitive tasks in which variables of interest are not directly observed and only information about their rate of changes is available (e.g., mental object rotation) [25, 75, 27]. MADE models could then act as test beds to generate mechanistic hypotheses for the network dynamics underpinning integration computation in such cognitive tasks.

### Activity bumps and tuning curves

MADE provides a basic prescription for the construction of continuous attractor and integrator networks of a desired dimension and topology. We numerically implemented a particular (Gaussian) kernel shape to illustrate the framework. The shape of the population activity bumps that result will depend on the kernel shape, which can be varied and selected as desired, according to the constraints supplied by our theory. Recent theoretical work on symmetry breaking for pattern formation also suggests that the set of potential kernels forms a large function space.

The tuning curve shapes of single cells depends both on the population activity bump shape as well as on the mapping from the external variable manifold to the internal neural state space manifold. As we have seen, if the external manifold is unbounded in some dimension but the internal representation is compact and periodic, then the spatial tuning curve will be periodic in that dimension. More subtle details of the mapping can affect the geometry of the periodic mapping, as we have described above.

We have focused our illustrations on simple and non-trivial manifolds of intrinsic dimension ≤ 2 for visualization and convenience. However, the theory and recipe for continuous attractor and integrator network construction generalizes in a straightforward manner to manifolds of higher dimension and different topologies.

### Related work

Computational models first described attractor networks [76, 36, 34, 35, 15, 11] and the mechanisms by which they could enable velocity integration [34, 47, 35, 15, 11, 46, 48, 18] long before experimental data verified the existence of such mechanisms. Intriguingly and surprisingly, in every case experimentally probed to date, the proposed neural circuit models closely resemble the hand-designed attractor models [40, 3, 42, 43, 44, 38, 15, 37, 14, 13]. Why is this the case? Presumably this match arises because the models were minimal in the sense that they implemented the essential elements and symmetries required to form the desired attractor, and circuits in the brain evolving under efficiency pressures arrived at similarly minimal models. MADE adopts a very similar mathematically minimal approach, recovering all of the known integrator models with bump-like tuning (except for the oculomotor integrator, which does not have bump-like responses).

An alternative approach to building models of integrating circuits in brains is to train artificial neural networks to perform tasks requiring integration [50, 49, 51]. After training, the networks’ solution is analyzed to reverse engineer the relation between network connectivity, neural dynamics and task performance [50, 77, 78, 79]. However, such approaches often fail to provide novel testable predictions or interpretable mechanisms to guide further experimental investigations, unless there was already a hand-crafted model available to which the trained network could be compared.

The network engineering approach [80, 81, 82, 83, 84, 82, 85, 86, 87] constructs circuits starting from the detailed desired dynamics of a system (precise states, fixed points, or specific tuning curves), then directly searching or solving for some network connectivity with those dynamics. Typically, these works further constrain the problem to make it well-posed by searching for low-rank weights or the lowest-dimensional embedding space for the dynamics while satisfying the desired properties. These methods are complementary to our approach: they permit construction of a broader set of dynamical systems, for instance trajectories ending in discrete fixed points, stable and unstable fixed points, etc., while our focus is specifically on biologically plausible continuous attractors that integrate. Conversely, those approaches do not provide a framework for building biologically realistic continuous attractor networks that integrate and lack known matches or easy interpretability to compare with biological circuits in known cases.

In conclusion, MADE allows for easy generation of interpretable, mechanistic, models of CAN networks that can integrate. We hope that MADE will endow researches with tools required to generate detailed, testable, hypotheses about the neural underpinnings of integration in diverse settings and in various cognitive processes, accelerating our understanding of the critical role that this class of computations play in many aspects of brain function and allowing for easy incorporation of such circuits in deep learning applications.

## Methods

All simulations and figures were implemented in custom Julia code available at GeneralAttractorsTheory. We will provide a minimal Python package for creating CANs and QANs using MADE: MADE-Python.

### CAN construction

In MADE, CAN engineering depends on computations of the pair-wise on-manifold distances between neurons in a lattice 𝒫. Thus, we begin by specifying a set of *n* equally spaced points on 𝒫. For the Line attractor, *n* = 256 and 𝒫 was taken to be the interval [−6, 6]. For the Ring attractor, *n* = 256 and 𝒫 was taken to be the interval [0, 2*π*] with the two ends identified (i.e. we ensured not to have a neuron at *θ*_*i*_ = 0 and one at *θ*_*i*_ = 2*π*). For all remaining networks, *n* = 48^2^ was used. The following rectangular intervals were used: for the plane attractor 𝒫 = [−10, 10] × [−10, 10], cylinder: 𝒫 = [−5, 5] × [0, 2*π*], torus: 𝒫[0, 2*π*] × [0, 2*π*], Möbius band: 𝒫 = [−2, 2] × [0, 2*π*] and Klein Bottle: 𝒫[0, 2*π*] × [0, 2*π*]. For the sphere attractor, the *n* points were chosen to be on a Fibonacci spiral on the unit sphere embedded in ℝ^3^.

Next, to implement custom manifold-specific distance metrics *d* we used the Julia package Distances.jl. The standard Euclidean metric was used for the line and plane attractor, for the ring a one dimensional periodic Euclidean metric (period 2*π*) was used, for the torus a two dimensional periodic Euclidean metric (period 2*π* in each direction) and for the Cylinder a heterogeneous periodic and standard Euclidean metric for the periodic and non-periodic dimensions respectively. For the sphere the great arc spherical distance for points on the unit sphere (implemented in the Manifolds.jl package [88]) was used. For the Möbius band a custom metric function was used to account for the non-orientable nature of the manifold.

For the Klein Bottle, a different approach was used. First, we defined an embedding of the Klein Bottle in ℝ^4^ mapping each lattice point *θ* = (*u, v*) to a point *q* ∈ ℝ^4^:

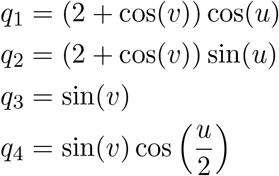

Next, we computed the pairwise Euclidean Distance in ℝ^4^ for the embedded points and selected the 8 nearest neighbors of each point. We then constructed a graph where each node was a lattice point and two nodes were connected if one belong to the neighborhood of the other. Each edge was assigned a weight equal to the Euclidean distance between the two points. Thus, the graph structure was taken to represent the local topological structure (connectivity) of the Klein Bottle. Given two points *θ*_*i*_, *θ*_*j*_ then, their on-manifold distance was given by summing the edge weights (local distances) along the shortest path on the graph from the node corresponding to *θ*_*i*_ to the one corresponding to *θ*_*j*_ as a way to numerically approximate the geodesic distance between them.

Following computation of pairwise distances, the connection weights between two neurons was computed using as kernel function:

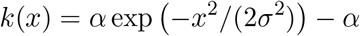

yielding strictly non-positive values for the connection strength. This gave a connectivity pattern characterized by global, long-distance inhibition, and no, or reduced, inhibition locally such that a localized pattern of activation on the neural lattice 𝒫 would remain localized and not result in activation of all neurons in the network. The parameters *α, σ* were varied based on the CAN topology and are indicated in table 1

**Table 1:** Kernel function parameters.

|  | line | ring | plane | cylinder | torus | Möbius | Sphere | Klein Bottle |
| --- | --- | --- | --- | --- | --- | --- | --- | --- |
| $\alpha$ | 1 | 1 | 2.5 | 2.5 | 2.5 | 2.5 | 2.5 | 2.5 |
| $\sigma$ | 1 | 1 | 25 | 25 | 2 | 2.5 | 40.5 | 150 |

### CAN simulation

Network dynamics were approximate to discrete time using forward Euler integration with Δ*t* = 0.5ms using:

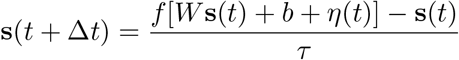

where **s**(*t*) is a vector representing the activity of each neuron in the network at time *t, τ* = 5*ms* was used as time constant. The constant input *b* = 0.5 was used throughout. The term *η*(*t*) was used to simulate Poisson noise in the recurrent dynamics, it represents a vector of length *n* whose entries are given by: 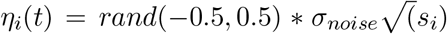 where *σ*_no*i*se_ ∈ {0, 1.5, 3, 5}. Unless explicitly stated, *σ*_no*i*se_ = 0 was used.

For each CAN, 2500 simulations of 25ms in duration were performed to generate data for the analysis of the activity manifold topology. We chose 25ms since we observed this to be sufficient for the network to settle into a steady state (i.e. one in which the network’s activity does not change between simulation steps).

For the first 15ms of each simulation, the activity of neurons at a distance *d >* 0.5 from a selected neuron *θ*_0_ (randomly selected for each simulation) was artificially set to 0 to induce the formation of a stable bump of activity around *θ*_0_ to promote uniform coverage of the entire manifold. The final activation vector **s**(*T*) for each simulation was then stored for subsequent analysis. For the torus attractor network, additional simulations were performed varying the noise parameter to assess the effect of noise on the attractor dynamics.

### Attractor manifold analysis

The final activation vector of each of 2500 CAN simulations for each manifold were collected into a matrix of dimensionality *n* × 2500 with *n* being the number of neurons in the network. For networks other than the line and ring attractors in which *n >* 400 a first dimensionality reduction step using PCA was performed to reduce the data to a 400 × 2500 dimensional matrix. Then, further reduction to three dimensional data for visualization (Figure 4) was achieved using Isomap [89]. To reduce computation Isomap was fitted to 10% randomly selected data points and then used to embed the entire dataset for visualization. For subsequent Topological Data Analysis (TDA) point cloud data was subjected to PCA dimensionality reduction to generate a 200 × 2500 data matrix and Isomap was then used to further reduce dimensionality to 10 (Isomap fitted to 10% of the data) [3].

### Topological data analysis

To perform persistent homology analysis the Julia packages Ripserer.jl and PersistenceDiagrams.jl [90] were used. To reduce computation, the TDA filtration was computed using a subset of randomly selected data points (20% of the entire dataset) to obtain the persistence diagrams shown in Figure 4 A. Only intervals with a lifespan *>* 7 were kept to remove features due to noise and the number of persistent intervals of each dimension (up to two dimensional cavities) were counted to obtain Betti numbers, which were then compared with those expected for manifolds of the given topology.

### Visualizing neural tuning curves

To visualize neural activation turning curves over 𝒩 in Figure 4, we used PCA and ISOMAP to reduce the dimensionality of neural activity to three dimensions. We thus obtained 2500 low dimensional points which we colored according to the activity of one selected neuron in the corresponding neural state. To visualize activity over the neural lattice 𝒫, we started by selecting one random neural state from the 2500 simulations. Then, we uniformly sampled 𝒫 and for each location *θ*_*i*_ ∈ 𝒫 we identified the closest neuron in the CAN (by coordinates). We then colored each point in 𝒫 according to the activation of the closest neuron.

### Intrinsic manifold dimensionality analysis

To estimate the manifold’s intrinsic dimensionality all data points in the *n*-dimensional state space were utilized. Pairwise Euclidean distance between each data point was computed to obtain each data point’s *k* nearest neighbors (using the NearestNeighbors.jl package). While Euclidean distance in state space does not necessary match on-manifold geodesic distance on 𝒩 in general, on a sufficiently small scale a manifold’s Euclidean structure makes this approximation acceptable. Next, 250 random data points (10% of the total) were selected for estimation of local dimensionality in their neighborhood. For each, the *k* closest points were selected and PCA fitted to the data. The number *d* of principal components required to explain at least 75% of the data was used as estimate of local dimensionality and the manifold’s intrinsic dimensionality was taken to be the average across repeats. Thus, the dimensionality estimation procedure depended on two hyperparameters: *k* and the percentage of variance explained. Preliminary tests on artificially generated data with known dimensionality and variable Gaussian noise were used to select the parameters used here, and we’ve found the estimated intrinsic dimensionality to be robust across a wide range of parameters values (data not shown). For the analyses shown here we used *k* = 500 throughout. Our preliminary tests showed that much smaller values of *k* resulted in noisy estimates (especially in the face of noise) and very large values of *k* led to an overestimation of the manifold intrinsic dimensionality (likely due to the higher global embedding dimensionality).

### Attractor dynamics analysis

To explicitly quantify attractor dynamics, a torus network was constructed as described above and simulated without external stimuli for a simulation time of 250ms (given 100 random initializations). Next, the network’s state was perturbed by addition with a random vector *v* of the same dimensionality as the network activity. For each of 100 simulations the vector was chosen to have a random orientation but fixed magnitude. The magnitude was computed to be 50% of the average distance between states on the torus manifold and the origin. Following the stimulus, the simulation was continued for 750ms more for a total simulation time of 1000ms. Data for each simulation was collected for the analysis of on-vs off-manifold dynamics. For each repeat the state just prior to stimulus application was used as seed for local PCA using k-nearest points from the point cloud data used for previous estimation of manifold topology, as described above (i.e. the steady states from previous simulations without the inputs were used to estimate the tangent plane to the manifold). The top two principal components were retained as approximation of the manifold’s tangent plane. The neural trajectory was then decomposed into on-manifold and off-manifold components by projection onto the tangent plane and remaining *N* − 2 dimensions. The euclidean distance in each subspace from the initial condition over time was then computed to asses drift following stimulus application and averaged across repeats.

### QAN construction

To construct quasi-periodic attractor networks for integration, first a choice of variable (ℳ) and neural lattice (𝒫) manifolds was made, ensuring that an identity or periodic mapping existed between the two (unless explicitly stated otherwise). Next, a map *π* : ℳ → 𝒫 and its inverse *π*^−1^ were defined (e.g. mapping each point on the plane, 𝒫, to a corresponding point on the torus, 𝒩). To compute connection weights in each QAN, points on 𝒫 were selected as before and the same distance metrics and kernel functions were used applying an offset to the neurons’ coordinates during distance computation. For the Line and Ring attractor, two QANs were constructed using the offset vectors *δ*_±m_ = ±0.15. For the Plane, Cylinder, Torus, Möbius attractors four QANs were constructed using *δ*_±1_ = [±0.25, 0] and *δ*_±1_ = [0, ±0.25] as offset vectors. For the sphere attractor, six QANs were constructed using as offset vectors *δ*_1±_ = ±[0, −*z, y*], *δ*_2±_ = ±[*z*, 0, −*x*] and *δ*_3±_ = ±[−*y, x*, 0] where [*x, y, z*] represents coordinates on the unit sphere embedded in three dimensional euclidean space. For the sphere, therefore, the offset vector magnitude varied as a function of position on the sphere to ensure that Killing vector fields were used (which are constant for the other manifolds used). The same vectors were used to compute the velocity-dependent stimulus 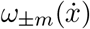 to each QAN. For some simulations, non-Killing vector fields where used. In the Torus, the offset vectors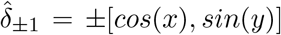, where *θ* = [*x, y*], and 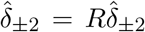 where used (where *R* is the rotation matrix *R* = [[0, 1], [−1, 0]]. For the sphere, the same vectors as above were used, except they were normalized to be of unit length everywhere on the sphere (except where they vanished).

### QAN dynamics simulation

Similarly to CANs, network dynamics were simulated using forward Euler integration. For each QAN in a network performing integration the discrete time dynamics were:

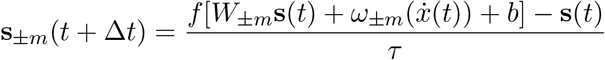

where **s**(*t*) =∑_±m_ **s**_±m_ and 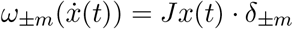 where *J* is the Jacobian of the map *π* evaluated at a point 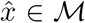 decoded from the neural state **s** and *δ*_±m_ is the offset vector at a point *θ* ∈ 𝒫 corresponding to the location of the neuron with highest activation in the network. The velocity input 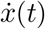 was computed by simulating the random walk of a point particle in the variable manifold ℳ.

To assess integration accuracy we generated 50 random trajectory (each corresponding to 1 second of simulated time) and simulated integration with the QANs. For each simulation, a trajectory 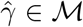 was decoded from neural activity and compared to the input trajectory *γ* ∈ ℳ. The simulation error was computed as a fraction of the trajectory length and was given by:

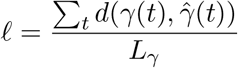

where *d* is a metric function for ℳ as described above and *L*_γ_ the trajectory length of *γ* on ℳ. More precisely, for decoding we used 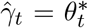 where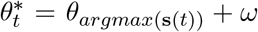. Here *ω* is a correction factor used only when ℳ and 𝒩 had different topologies such that ℳ had non-periodic dimensions (plane) and 𝒩 had periodic dimensions (torus). At each decoding step, we added or subtracted 2*π* to *ω* when necessary to account for the neural state “wrapping around” the boundary dimension(s). Here *ω* is a *d*-dimensional vector and each value is set to 0 for non-periodic dimensions in 𝒩 and is *k*2*π* for some integer *k* otherwise.

To generate the trajectories, we first defined a set of 2*d* vector fields Φ_*i*_ over ℳ, each corresponding to a vector field Ψ_*i*_ over 𝒫. Then, we generated smoothly varying vectors *A*^*i*^ such that at each time *t* the velocity vector 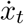 was given by 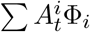. These weights vectors were given by the sum of two sine waves with random periods and scaled to have amplitude *<* 0.1. We then computed **X**, the trajectory over ℳ by, at each time step, computing 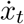 and 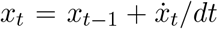(with *dt* the simulation time step, 0.5). Finally, we computed **V** the set of inputs to the QANs. For each time point *t*, the input *v*_*j*_ to the *j*^th^ QAN was given by 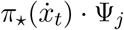 where *π*_⋆_ was the push forward of the map *π* : ℳ → 𝒩.

In some conditions we artificially injected Poisson noise in the QAN neural dynamics as described previously to assess the effect of noise on path integration.

### Neural tuning curves on ℳ

To visualize neural tuning curves with respect to ℳ in Figure 6, we generated a single trajectory densely sampling from ℳ (5-10 seconds of simulated time). After simulating path integration, we selected one random neuron to visualize its tuning curve. The visualization method varied based on the manifold topology. For one dimensional manifolds we simply plotted decoded value, *x*, against the neuron’s activity. For most two dimensional manifolds, with the exception of the sphere, we generated a heatmap by binning *x* and quantifying the average neuron’s activity for samples from each bin. A small amount of noise was added to *x* before binning to improve visualization. For the sphere, we first sampled 2000 points uniformly distributed on ℳ. Then, for each point we looked at the closest decoded value. We then colored each point on the sphere according to the neural activity value at the corresponding sample.

### Non Killing fields and non-periodic manifold mapping

To demonstrated that path integration depended on the weight offset vector fields Ψ being Killing fields we generated two variants of the torus and sphere QANs. For the torus, we kept the magnitude and relative orientation of the offset vector fields constant, but gradually rotated their position by an angle *cos*(*θ*_1_) (i.e. only as a function of position along one manifold dimension). This ensured that vector fields at the boundary conditions were identical, as expected. For the sphere, we started with the Killing vector fields we had, and simply normalized each vector such that all vectors had constant length. We then ran 50 simulations using random trajectories as described previously.

To assess path integration when no trivial or periodic mapping between ℳ and 𝒩 existed, we performed path integration simulations with ℳ as a cylinder and 𝒩 as a sphere. We used the same procedure described above to generate 50 random trajectories over the cylinder and computing the corresponding velocity vectors over 𝒫.

## Supplementary Information

**Figure S1:**
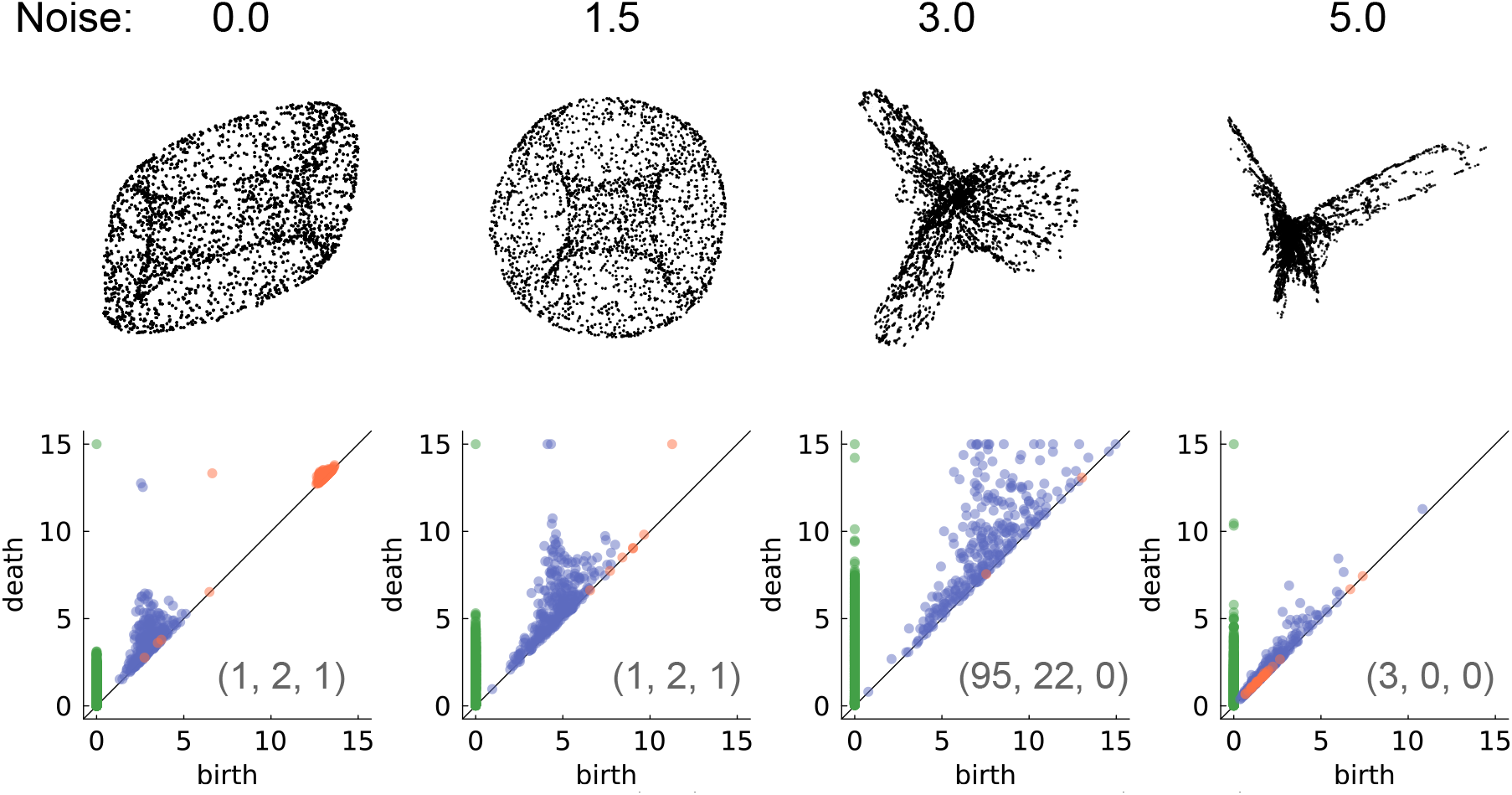
Torus CAN activity manifold (top) and persistence diagram (bottom) for varying noise intensity levels (columns).

### 1 Kernels constructed through distance metrics produce single bump states

Here we estimate the conditions on the interactions that lead to the formation of bump states on the lattice of neurons, 𝒫.

As earlier, consider an interaction weight matrix *W* (*θ, θ*^′^) = *k*(*d*(*θ, θ*^′^)). We rewrite the kernel *k* as *k*(*d*) = −*k*_0_ + *k*_1_(*d*), where *k*_1_(*d*) → 0 as *d* → *∞* and *k*_1_(0) = *k*_0_ *>* 0; and correspondingly write *W* (*d*(*θ, θ*^′^)) = −*W*_0_ + *W*_1_(*d*(*θ, θ*^′^)). We assume that the kernel *k* has a length scale *σ*, i.e., *k*_1_(*d*) *≈* 0 for *d* ≥ *σ*.

Since *σ* is the only spatial scale being introduced in the dynamics, we qualitatively expect the localized bump states will have a scale of O(*σ*). If *σ* is much smaller than the distances over which the manifold 𝒫 has curvature, 𝒫 will be approximately flat within a ball *V*_σ_ centered on any *x* ∈ 𝒫. In this approximation, the conditions for the formation of a stable bump state are the same as those for the formation on a bump state on a globally flat manifold.

To examine the conditions for the existence of a bump state, we will first calculate the homogeneous steady state supported by Eq. 1. Next, we note that since *W* is symmetric in this case, thus Eq. 1 can be described through an energy function[58], and thus a stable steady state must exist. If the homogeneous state is unstable, there must then exist a stable symmetry broken state of the system. If this symmetry broken state is localized, we refer to it as the bump state.

The homogeneous steady state *s*(*x*) = *s*_0_ must satisfy

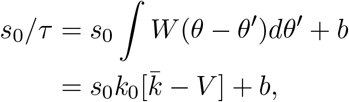

where *V* is the volume of the manifold, ∫*dθ*, and 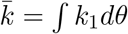. Rearranging, we obtain

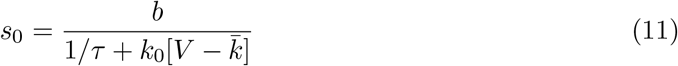

Since the kernel *k*_1_ is supported on a small volume of the entire manifold, 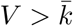, and thus the right-hand side of Eq. 11 is positive, consistent with the assumed rectifying nonlinearity *f* of Eq. 1.

To examine the stability of this homogeneous state, consider a small perturbation, *s*(*x, θ*) = *s*_0_ + exp(*α*(*ω*)*t* + *iω* · *θ*) to Eq. 1. Following the analysis in Ref. [59], we obtain

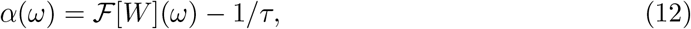

where ℱ [*W*] is the Fourier transform of the interaction *W* .

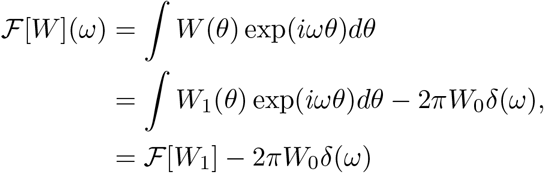

where *δ*(*ω*) is the Dirac delta function, obtained from the Fourier transform of a constant. Thus, the homogeneous steady state will be unstable if ℱ [*W*](*ω*) *>* 1*/τ* for some *ω*. Since *α*(*ω*) denotes the rate of exponential growth, the maxima of Eq. 12 will determine the dominant growing mode. If ℱ[*W*] were maximized at |*ω*| *>* 0, then the growing perturbation would have a periodic component, and would thus likely not form a localized mode. Instead, if ℱ[*W*_1_](*ω*) were maximized at *ω* = 0, then ℱ[*W*](*ω*) will be maximized at *ω* → 0 (ℱ[*W*](*ω*) cannot be maximized strictly at *ω* = 0 itself due to the −2*πW*_0_*δ*(*ω*) contribution to ℱ[*W*](*ω*)). In this case, the growing perturbation will be unimodal, likely leading to the formation of a localized state.

Thus, for the formation of a stable bump state on a general manifold, we obtain two requirements: First, the Fourier transform of the kernel *k*_1_(*d*) must be maximized at *ω* = 0; and second, this maximum must be larger than 1*/τ*. If we are solely interested in interaction shapes that lead to bump formation, we assume we have freedom to rescale the interactions. Thus, if a positive maximum is attained at *ω* = 0 a rescaling can always make this maximum larger than 1*/τ*. Thus, we primarily focus on the first requirement.

While we do not provide an exhaustive classification of interaction kernels *k*_1_ whose Fourier transforms are maximized at zero, we provide a broad sufficient condition — if *k*_1_(*d*) ≥ 0 for all *d*, then its Fourier transform will be maximized at zero. This can be proved as:

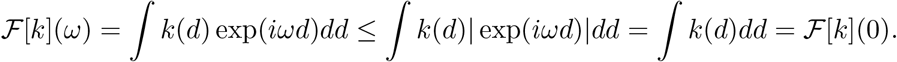

Thus, we finally conclude that, up to a rescaling of the strength of the interaction, an interaction *W* (*d*(*θ, θ*^′^)) will lead to the formation of a bump state if it can be rewritten as *W* (*d*(*θ, θ*^′^)) = *k*_1_(*d*(*θ, θ*^′^)) − *k*_0_ for: *k*_0_ ≥ 0; a kernel *k*_1_ that satisfies *k*_1_(*d*) ≥ 0 and *k*_1_(*d*) → 0 for *d* ≥ *σ* ; and sufficiently small *σ* over which the manifold 𝒫 is approximately flat.

### 2 Manifold of single bump states 𝒩 is isometric to manifold of neural lattice 𝒫

Here we will show that the manifold 𝒩 of neural activity, formed through single bump states at each point of the neural lattice 𝒫, is isometric to 𝒫. Specifically, we provide a distance metric *d*_*N*_ on the manifold 𝒩, such that (𝒩, *d*_*N*_) is isometric to (𝒫, *d*_*P*_), where *d*_*p*_ represents the geodesic distance considered as the distance metric during the MADE construction described in the main text.

While we will not prove this in complete generality for any 𝒫, we will assume that if 𝒫 has a sufficiently large separation of lengthscales (as assumed in the previous section), it will suffice to show this result for 𝒫 given as the flat Eucldiean manifold ℝ^n^ (and correspondingly, *d*_*p*_ being the usual *L*_2_ metric).

To prove the existence of an isometry, we first argue that 𝒩 and 𝒫 are diffeomorphic. In Apx. 1, we argued that the prescribed connectivity kernel leads to the formation of activity bump states centered at any *x* ∈ 𝒫. Define the function *f* from 𝒫 to 𝒩 to characterize the shape of the activity bump, i.e., for any *x*_0_ ∈ 𝒫, we let 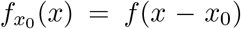 be the shape of the activity bump centered at *x*_0_. Since these activity bump states are generated through radially symmetric kernel interaction functions, the bump states *f* (*x* − *x*_0_) must also be radially symmetric, i.e., *f* (*x* − *x*_0_) = *F* (|*x* − *x*_0_|). In this case, we can see that Φ : *x* → *f*_x_ is now a diffeomorphism, since it is a smooth function and has a smooth inverse (the inverse map is simply computing the center of the radially symmetric activity bump).

Note that a direct *L*_2_ norm on 𝒩 does not suffice, since for sufficiently distant *x*_0_ and *x*_1_,

Next, we examine candidate metrics on 𝒩 that may lead to an isometry with (𝒫, *L*_2_). the distance between 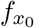 and 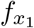 given by 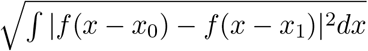 is approximately 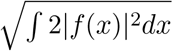. Thus the distance between 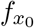 and 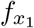 is bounded, whereas the distance between *x*_0_ and *x*_1_ is not, indicating that there cannot exist a direct isometry.

Instead, we construct here a metric of intrinsic length induced by the Riemannian metric on the tangents of 𝒩. For any two vectors *u*(*x*) and *v*(*x*) in *T*_s_ 𝒩, the tangent space of 𝒩 at *s*. Define the Riemannian metric as *g*(*u, v*) = ⟨*u, v*⟩ =∫ *uvdx*. Then, for any path *γ*(*t*) ∈ 𝒩, we can define the length of the path *L*[*γ*(*t*)] as

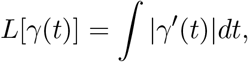

where the norm of a tangent vector *γ*^′^ is defined as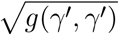. This can now be used to define the geodesic metric between 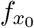 and 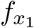 on 𝒩 given as the infimum of the lengths of all paths between 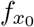 and 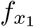. Here we will show that under this geodesic metric, the spaces (𝒩, *d*_*N*_) and (*P, d*_*P*_) are isometric. Specifically, we will show that the metric tensor (the Riemannian metric computed for coordinate basis vectors) is proportional to identity, the metric tensor for flat Euclidean space.

Assume that 𝒩 is an *n* dimensional manifold. Let (*x*^1^, …*x*^n^) be a coordinate chart in the neighborhood of a bump state 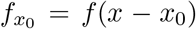. A basis for the tangent space is then given by the differentials {*∂/∂x*^1^, …*∂/∂x*^n^}. Note that since *f* (*x*) is radially symmetric, *f* (*x*) = *F* (|*x*|), the basis vectors can be simplified as

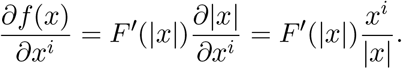

We can now compute the metric tensor *g*_*ij*_ = *g*(*∂/∂x*^*i*^, *∂/∂x*^j^)

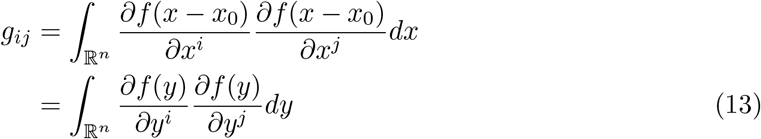

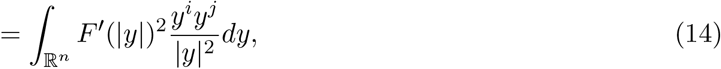

where Eq. 13 is obtained by performing the change of variables *y* = *x* − *x*_0_. From Eq. 14 we can make two crucial observations: first, since the integrand is odd in *y*^*i*^ and *y*^j^, thus *g*_*ij*_ = 0 for *i* ≠ *j*; second, *g*_*ii*_ is independent of *x*_0_, and by symmetry is also independent of *i* — it is entirely determined by the shape of the bump state *F* (|*x*|). Thus, the metric tensor *g*_*ij*_ has constant entries on the diagonal, and zero on the off-diagonal elements, i.e., *g* is proportional to the identity matrix. We denote this proportionality constant as *α*.

The length of an infinitesimal line element *ds* is then given as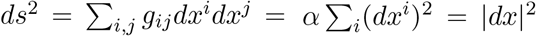. The length of a path *γ* from 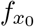 to 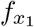 is then simply 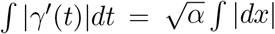, which is the Euclidean path length from *x*_0_ to *x*_1_ scaled by 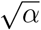. Thus, the geodesic metric from 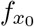 to 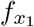 is the infimum of Euclidean path lengths, i.e., the Euclidean straight-line distance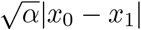. We can additionally redefine a new metric 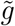 on the tangent space as *g/α*, leading to the new geodesic distance to be exactly the Euclidean distance |*x*_0_ − *x*_1_|.

Thus, under the approximation of 𝒫 being treated as a flat space without curvature at scales smaller than *σ*, the metric space (𝒩, *d*_*N*_) is thus isometric to the metric space (𝒫, *d*_*P*_).

### 3 External velocities ignorant about network structure and state require shifted-kernel networks to control bump flow

In this section, for analytical simplicity, we will ignore the neural transfer function nonlinearity *f*.

The fixed points resulting from symmetric kernels in Eq. 1 satisfy:

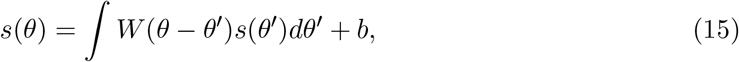

where *s*(*θ*) denotes an activity bump centered at any point in 𝒫. Consider two such activity bump states: *s*_0_(*θ*) centered at *θ*_0_, and a nearby state *s*_ϵ_ centered at *θ*_0_−*ϵ*, i.e., *s*_ϵ_(*θ*) = *s*_0_(*θ*+*ϵ*). For a neural state *s*(*θ*) to move from *s*_0_ to *s*_ϵ_ in time Δ*t*, the time derivative *∂s/∂t* must equal

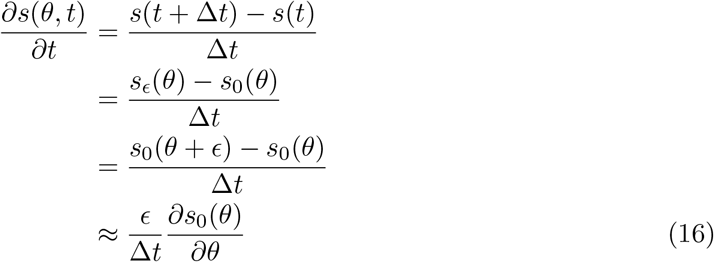

We can use Eq. 15 to evaluate this space derivative as

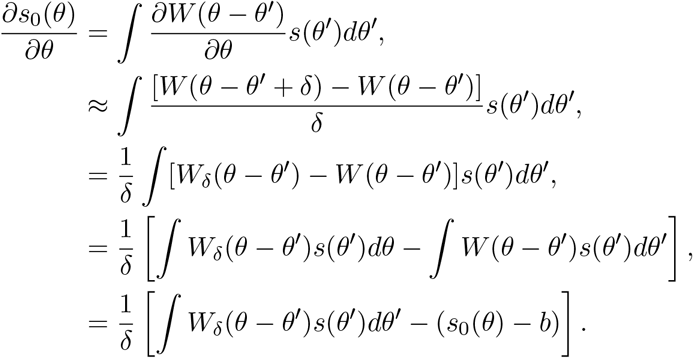

where *W*_*δ*_ represents a kernel with a small offset *δ*, i.e., *W*_*δ*_ = *W* (*θ* − *θ*^′^ − *δ*). We can insert this in Eq. 16 to obtain

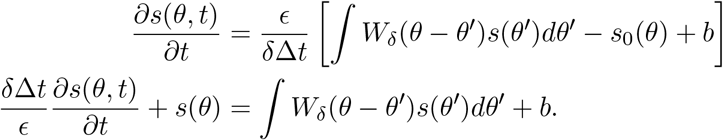

Comparing the above equation with Eq. 1, we find that the neural time constant *τ* = *δ*Δ*t/ϵ*. Since the speed of the activity bump is *ϵ/*Δ*t*, we obtain a speed of

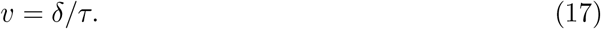

Thus, a network built with a kernel with offset *δ* in particular direction leads to activity flow along that direction. Coupling multiple copies of such networks with opposing directions of kernel offsets leads to an equilibrium, with the bump state at a fixed position. This can be intuitively seen by noting that *W*_*δ*_*s* + *W*_−*δ*_*s ≈* (*W* + *δ∂*_θ_*W*)*s* + (*W* − *δ∂*_θ_*W*)*s* = 2*Ws*, and thus opposing offset kernels acting on the same state are equivalent to the state being acted on by a kernel with no offset.

To control the flow the bump in arbitrary directions, we will next demonstrate that the magnitude of the feed-forward input *b* in a particular subnetwork can bias the motion of the bump. To see this, we first consider Eq. 1 scaled by a factor *α*,

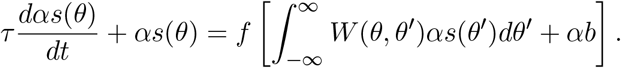

Thus, scaling *b* by a factor *α* (i.e., *b* → *αb*) results in an equivalent solution of the dynamical equation with the states *s* also scaled by the same factor *α* (i.e., *s*(*θ*) → *αs*(*θ*)).

Consider two such coupled networks with opposing offsets, with feedforward inputs scaled by *α*_1_ = (1 + *α*)/2 and *α*_2_ = (1 − *α*)*/*2. As noted above the neural firing rates can be assumed to be scaled by the same factors. Heuristically, we will assume that the firing rates of the coupled network can be approximated through individually scaled firing rates of independent offset networks. This leads to the effective interaction through the offset kernels as

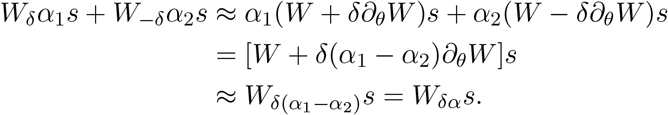

Thus, the effective interaction is similar to that obtained by a kernel with an offset of *δα*, leading to a bump speed of *δα/τ*.

Finally, we note that while the above argument has been constructed for offsets along a single dimension, it readily generalizes to higher dimensions: For continuous and differentiable *W*, a directional derivative can be written as a linear combination of partial derivatives along coordinate axes, i.e.,

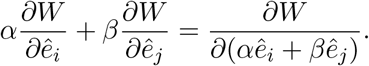

Thus, subnetworks with differently scaled feedforward inputs lead to differently scaled firing rates *s* which leads to an interaction kernel that has an effective offset in the vector direction determined by the scaling coefficients. This effective offset in a particular direction causes the activity bump to flow along the manifold along that direction, leading to controllable flow of the activity bump through differential feed-forward inputs to the coupled network.

